# Re-ranking of Computational Protein–Peptide Docking Solutions With Amino Acid Profiles of Rigid-Body Docking Results

**DOI:** 10.1101/2020.05.12.092007

**Authors:** Masahito Ohue

## Abstract

Protein–peptide interactions, in which one partner is a globular protein and the other is a flexible linear peptide, are important for understanding cellular processes and regulatory pathways, and are therefore targets for drug discovery. In this study, I combined rigid-body protein-protein docking software (MEGADOCK) and global flexible protein–peptide docking software (CABS-dock) to establish a re-ranking method with amino acid contact profiles using rigid-body sampling decoys. I demonstrate that the correct complex structure cannot be predicted (< 10 Å peptide RMSD) using the current version of CABS-dock alone. However, my newly proposed re-ranking method based on the amino acid contact profile using rigid-body search results (designated the decoy profile) demonstrated the possibility of improvement of predictions. Adoption of my proposed method along with continuous efforts for effective computational modeling of protein–peptide interactions can provide useful information to understand complex biological processes in molecular detail and modulate protein-protein interactions in disease treatment.

## 1 Introduction

Protein–peptide interactions involving a globular protein and a flexible linear peptide are important for understanding cellular processes and regulatory pathways; hence, such interactions are common targets for drug discovery [1]. Many recent studies reported the biological functions of short open reading frames-encoded peptides or micropeptides encoded in noncoding RNAs [2, 3, 4, 5]. However, many peptide- and protein–peptide interactions remain to be uncovered, which can drastically shift the field of traditional genome biology.

Various protein–peptide docking prediction tools have been developed recently for determination of the structures of protein–peptide complexes [6, 7, 8, 9, 10, 11] and reviewed in [12]. Protein–peptide docking involves certain computation steps such as initial conformation sampling, structural refinement, and rescoring, similar to traditional protein-protein docking or protein-ligand docking techniques. Thus, these prediction tools offer broad steps that can be used for a single purpose. CABS-dock [10, 11] is one of the few tools that is capable of predicting a protein–peptide complex in a one-step process based on protein structure and peptide sequence information. Specifically, CABS-dock generates candidate complex structures with C*α*-C*β*-side group protein model (CABS model [13])-based replica exchange Monte Carlo dynamics simulations and then performs all-atom modeling by MODELLER [14]. With the rapid increase in the generation and deposition of peptide structure information in public database such as PeptiDB [15], template-based prediction is also a powerful tool for protein–peptide docking when the peptide template is identified [6].

Despite the wide array of methods recently developed in the field of protein-protein and ligand-peptide docking, it remains a challenge to effectively achieve protein–peptide docking. For example, the targets of Critical Assessment of PRedicted Interactions (CAPRI), T65, T66, and T67, are very difficult to predict accurately as protein–peptide complexes, resulting in frequent incorrect answers, especially for T65 and T66 [16]. Thus, further attempts at accuracy improvement for such predictions are very important. Despite apparent similarities, compared with the diverse prediction approaches available for protein-protein docking or protein-ligand docking, the field of protein–peptide docking methodology is still in an immature phase[17, 18].

The computational techniques used in protein-protein docking are arguably more important for protein–peptide docking. This is not simply because the peptide chain consists of amino acids (as in a protein), but several protein structure informatics can also be appropriated for protein–peptide docking. Protein-protein docking first developed with a fast Fourier transform-based exhaustive search technique using rigid bodies as one of the elemental technologies; MEGADOCK [19] is an example based this approach. By considering the structure of a protein as rigid, structural sampling can be achieved over the entire space with a short calculation time. The latest tools (MEGADOCK 4.0 [20] and Hex [21]) are capable of evaluating trillions of conformations in seconds on GPU accelerators, and protein-protein docking tools are used casually. Although the conformations predicted with these tools tend to include many false-positive complexes, the so-called decoy conformations that result from a rigid-body search can be used in reevaluating the predicted conformations [22, 23, 24] or provide information for the search direction in redocking [25]. This can expand the application of these tools from predicting a single complex structure to prediction of the post-docking processes.

However, rigid-body sampling is not sufficiently efficient to be used for highly flexible proteins [26]. This is a reasonable outcome considering the basis of “rigid”. Therefore, when dealing with flexible proteins, many more false-positive candidates will be generated. Accordingly, this approach has been considered to be difficult to apply to problems such as protein–peptide docking. In this study, I address this problem by incorporating the rigid-body sampling results to the re-ranking of protein–peptide docking solutions. Because sampling with a rigid-body model results in many false-positives, I propose the “decoy profile” as the environment in which the binding site is favored by rigid-body samples. I then developed a new protein–peptide docking re-ranking method with the decoy profile. The method can be performed with very light computation requirements. Moreover, since only rigid-body docking sampling is used, the method does not need the training datasets of protein–peptide complex structures that are required with machine learning-based methods.

## 2 Materials and Methods

### 2.1 protein–peptide Docking Re-ranking

An overview of the proposed method is presented in Figure 1. The purpose of protein– peptide docking is to predict the complex structure based on an input protein tertiary structure and a peptide sequence.

**Fig. 1.**
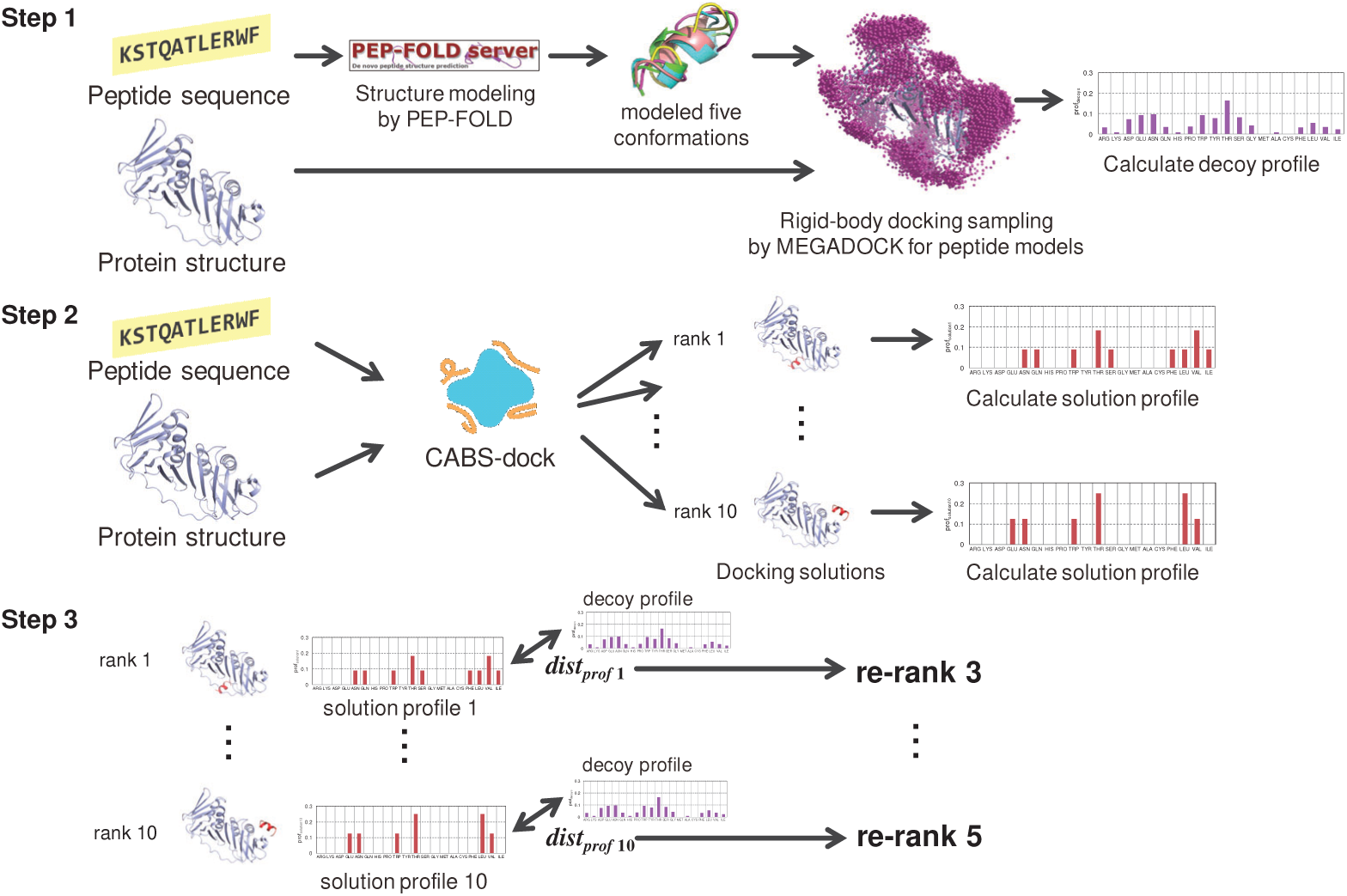
Overview of the proposed method.

In the proposed method, a vector of the relative frequencies of the number of protein residues with which the peptide makes contact (solution profile) is obtained for the generated protein–peptide complex solution. In addition, I independently obtain the residue contact profile of the decoys (decoy profile) among a large number of rigid-body docking samples to provide a clue for estimating the peptide binding preference. In other words, the decoy profile attempts to pseudo-estimate the ideal interaction surface environment for peptide binding to the target protein. Finally, the similarity between the solution profile and the decoy profile is determined by the Euclidean distance. A better solution is considered as one in which the solution profile is the closest to the decoy profile, and then the re-ranking process is performed accordingly.

The proposed method is conducted by the following three steps:

Step 1: calculate the decoy profile,
Step 2: calculate the solution profile,
Step 3: calculate the profile-profile distance and re-rank.

#### Step 1: Calculating the Decoy Profile

I generated five peptide conformations from the input peptide sequence using the PEP-FOLD server [27]. The input protein structure and modeled peptide structures were then docked using MEGADOCK rigid-body docking software [20] to sample numerous peptide docking poses simultaneously. Specifically, I sampled 3,600 poses for each peptide model, and a total of 3,600×5 = 18,000 poses were generated using MEGADOCK. The contact amino acid profile was then determined using these “decoy” structures (decoy profile) according to the following equation:

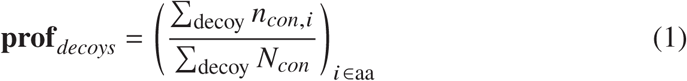

where *n*_*con,i*_ is the number of residues *i* in contact with the peptide, *N*_*con*_ is the total number of residues in contact with the peptide, and *i* ∈ aa represents the 20 standard amino acids. The judgment of residue-peptide contact is made when the distance between any of the heavy atoms of the protein residue *i* and any of the heavy atoms of the peptide is less than 4 Å. The values of *n*_*con,i*_ and *N*_*con*_ are then incremented.

#### Step 2: Calculating the Solution Profile

To obtain solutions of the protein–peptide complex, the following vector (solution profile) was calculated:

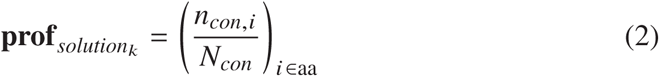

where *n*_*con,i*_ and *N*_*con*_ are defined as above for the decoy profile calculation.

Although some residues can more easily make contact, whereas others will be less able to make contact due to the difference in the number of amino acids in a protein, normalization by the relative frequency of amino acids in a protein was not necessary in this case because the profiles were only compared for the same protein–peptide target.

In this study, the solutions of the protein–peptide complex were generated by CABS-dock [11]. I ran CABS-dock with default settings using the target peptide sequence and protein structure, and then calculated the profiles of each of the 10 solutions generated by CABS-dock.

#### Step 3: Calculating the Profile-Profile Distance and Re-ranking

The profile-profile distance *dist*_*prof*_ between the decoy profile and each solution profile was calculated as follows:

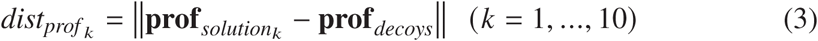

The solutions with smaller 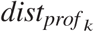 values were then re-ranked so that they come out on top.

### 2.2 Dataset

#### 2.2.1 protein–peptide Complex Dataset

Among the 10 protein–peptide complexes in the dataset generated by Dagliyan et al. [9], those with 9 or more residues were selected for use in this study. Tables 1 and 2 show the details of the dataset. The number of peptide residues ranged from 9 to 13; all hydrogen atoms and HETATM records in the PDB structure data were deleted. Only 1PRM was an nuclear magnetic resonance (NMR) structure. For the NMR structure, only the coordinates of the first state were used for the input structure.

**Table 1.** protein–peptide complexes.

| PDB ID | Protein | Peptide |
| --- | --- | --- |
| 1PRM | c-Src tyrosine kinase SH3 domain | proline-rich ligand PLR1 |
| 1X2R | kelch-like ECH-associated protein 1 | Nrf2/Neh2 peptide |
| 1RXZ | DNA polymerase sliding clamp | PCNA-binding motif |
| 2FMF | Chemotaxis protein CheY | C-terminal 15-mer helix of CheZ |

**Table 2.**
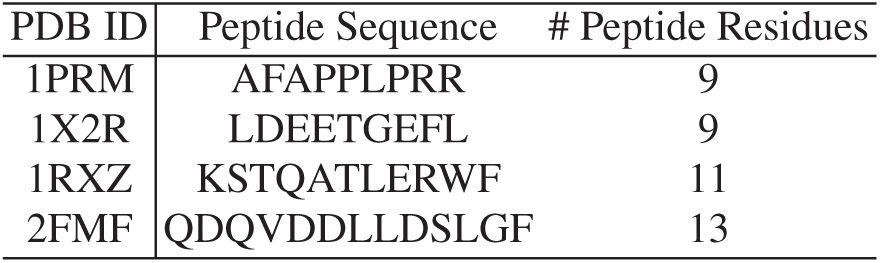
Peptide sequences.

#### 2.2.2 protein–peptide Docking Solutions

The holo (bound) state of the protein structure was obtained as a PDB file and the sequence information of the peptide was input to CABS-dock to obtain the docking solutions. The peptide secondary structure information was not used and the number of simulation cycles was set to 50, which is the default parameter for CABS-dock.

#### 2.2.3 Rigid-Body Docking Decoys

The sequence information of each peptide was input into PEP-FOLD, resulting in five peptide conformations for each peptide. The target protein structure and peptide conformations were then entered into MEGADOCK version 4.0.2 to obtain 3,600 rigid-body docking solutions (decoys) per peptide conformation for a total of 18,000 decoys. The protein PDB file was set to `-R` and the peptide PDB file was set to `–L` using the `-N 3600` option; default parameters were used for the other settings of MEGADOCK.

### 2.3 Evaluation of Prediction Performance

#### 2.3.1 Evaluation of the Predicted Structure

The docking solution was evaluated using the all-heavy-atom root mean square deviation (RMSD) between the native peptide structure and the docking solution peptide when the protein structure was superimposed. I defined a near-native solution (correct answer) when the RMSD was less than 10 Å. The RMSD was calculated using ProDy library [29].

#### 2.3.2 Solution Ranking Evaluation

The rankings of the 10 predicted structures output by CABS-dock with those obtained after re-ranking were compared using the proposed method. The area under the receiver operating characteristics curve (AUC), calculated from the rank position of the near-native solution (Eq. 4), was also used to compare the rankings as follows:

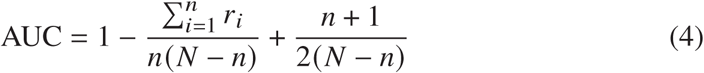

where *n* is the number of near-native solutions, *N* is the number of all solutions (= 10), and *r*_*i*_ is the rank of each near-native solution. For example, if the 2nd- and 4th-ranked docking solutions were found to be near-native among the 10 solutions, then *n* = 2, *N* = 10, *r*_1_ = 2, and *r*_2_ = 4; thus, 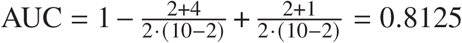. Note that in this study, the focus was on the top 10 solutions of CABS-dock, and the AUC value for 10 solutions will be different from the measure used for common docking results such as those dealing with thousands of candidates. For example, a result of *n* = 1, *N* = 10, *r*_1_ = 8 that yields a low value of AUC = 0.222 is considered to be a generally good result for protein–peptide docking. In this study, the AUC value was simply used as an indicator to check whether the ranking had been improved by my method.

## 3 Results and Discussion

### 3.1 CABS-dock and Re-ranking Results

Table 3 shows the RMSD, rank, and ranking change based on the proposed method, along with the AUC values of the docking solutions for CABS-dock. Protein–peptide docking with CABS-dock generated near-native solutions in the top rank with the exception of 1PRM.

**Table 3.** Results of CABS-dock solutions and re-ranking by the proposed method. Bold values represent near-native solutions.

| rank | 1PRM |  | 1X2R |  | 1RXZ |  | 2FMF |  |
| --- | --- | --- | --- | --- | --- | --- | --- | --- |
|  | RMSD (Å) | re-rank | RMSD (Å) | re-rank | RMSD (Å) | re-rank | RMSD (Å) | re-rank |
| 1 | 24.40 | 8 | **9.83** | 6 | **5.38** | 3 | **7.80** | 3 |
| 2 | 19.79 | 7 | **8.21** | 5 | **5.86** | 2 | **7.74** | 2 |
| 3 | 17.70 | 2 | 31.97 | 10 | 25.53 | 6 | **8.55** | 1 |
| 4 | 14.68 | 10 | 39.26 | 3 | 26.95 | 8 | **8.30** | 4 |
| 5 | 19.31 | 3 | 20.35 | 7 | 24.75 | 4 | 20.91 | 7 |
| 6 | 16.71 | 9 | 37.43 | 9 | 33.30 | 7 | 12.91 | 5 |
| 7 | 16.06 | 1 | **7.22** | 1 | 24.83 | 1 | 33.18 | 10 |
| 8 | **6.54** | 6 | **9.00** | 2 | 25.58 | 9 | **8.93** | 9 |
| 9 | 18.15 | 4 | **8.81** | 4 | 25.75 | 10 | 33.38 | 6 |
| 10 | 23.76 | 5 | **8.45** | 8 | 35.09 | 5 | 28.21 | 8 |
| AUC | 0.222 | 0.444 | 0.333 | 0.792 | 1.000 | 0.875 | 0.880 | 0.840 |

In addition, when considering the best near-native solution rankings for re-ranks, 1RXZ moved down a rank from first to second, but otherwise did not deteriorate. Overall, 1PRM, which had no near-native solution at #1, had the best near-native solution rank improvement by moving up from position 8 to position 6.

With respect to the ranking of near-native solutions based on AUC values, 1RXZ and 2FMF showed a slight deterioration in ranking, whereas 1PRM and 1×2R showed good improvement. In particular, the enrichment of near-native solutions for 1×2R improved significantly. The improved peptides were all 9-mer, suggesting that the effect of re-ranking may be higher when the number of residues is less than 10. However, this may also be related to the performance of the PEP-FOLD conformation sampling. Figure 2 shows the sampling results of PEP-FOLD. For the most successful case (1×2R), the conformation sampling of peptides by PEP-FOLD yielded a structure close to that of the native form. Conversely, no peptide conformation similar to the native form was obtained for 1PRM. Thus, it is likely that the performance of conformation sampling affects the re-ranking according to the decoy profile.

**Fig. 2.**
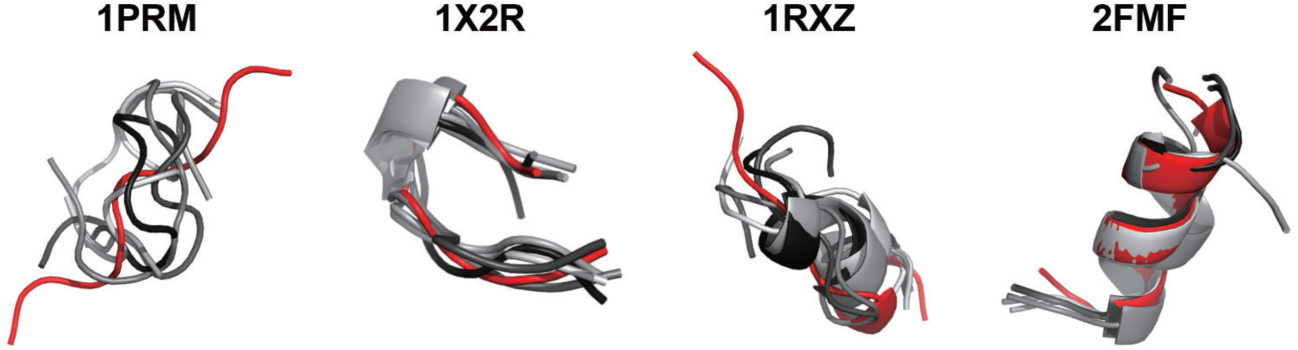
PEP-FOLD conformation sampling results. Red peptides represent the binding form obtained from the PDB complex and gray peptides represent the results from PEP-FOLD sampling.

### 3.2 Example of Predicted Structures

The native structure and some solutions for 1×2R are shown in Figure 3. As shown in Figure 3(a), we can confirm that the solutions located around the correct binding site have RMSDs less than 10 Å. Similarly, solutions with RMSDs greater than 10 Å bound at completely different sites than the correct site. The green-colored peptide in Figure 3, which ranked at the top after re-ranking, was at a position and orientation closer to those of the native-like form compared to the structure of the original top-ranked peptide (yellow) (Figure 3(b)).

**Fig. 3.**
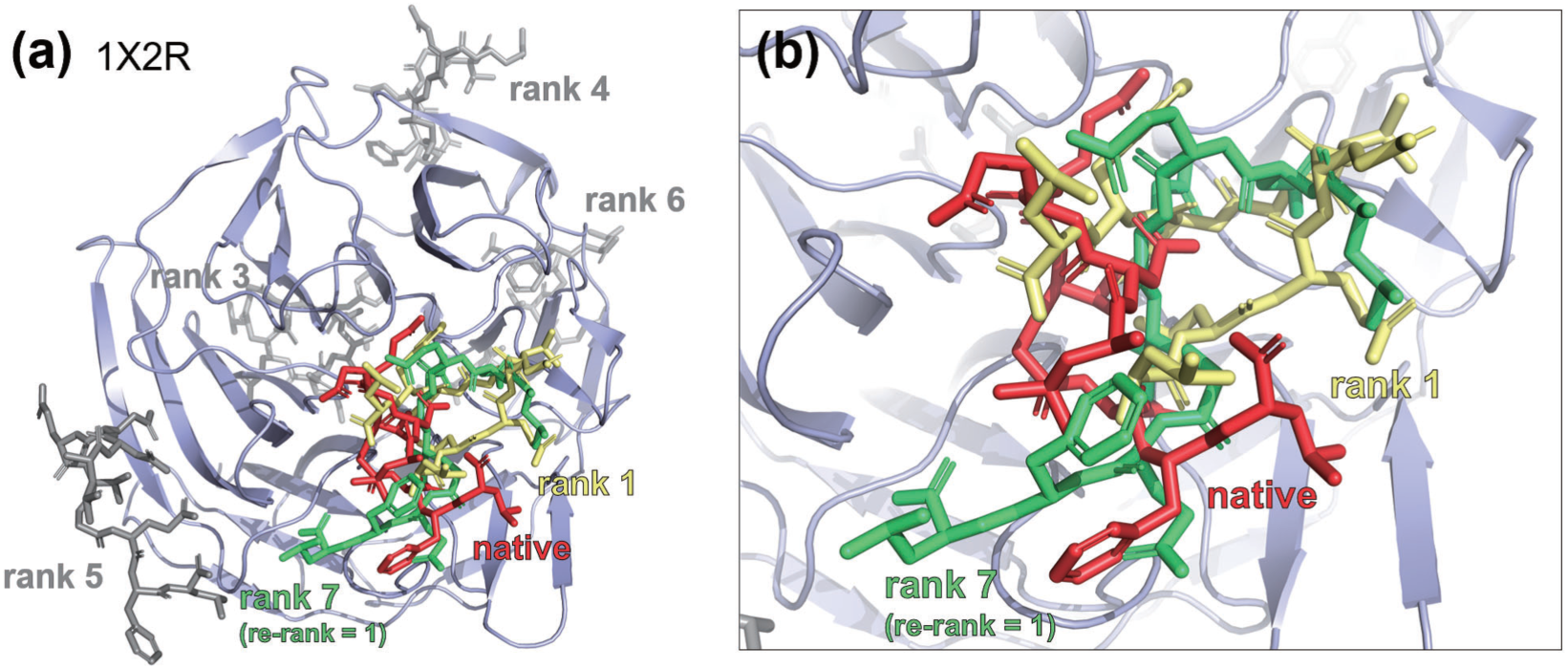
Results of 1X2R peptide docking and re-ranking. (a) Overall image of the protein, and (b) enlarged image of the area around the native binding site.

## 4 Conclusion

I have proposed a re-ranking method using amino acid residue contact information from rigid-body docking results to improve protein–peptide docking calculations. Application of the proposed method to four protein–peptide docking solution sets showed general improvement in the ranking of near-native solutions with my re-ranking approach. As the number of peptide residues increased, the degrees of freedom increased and improvement became generally more difficult; however, the re-ranking prevented worsening predictions.

Peptides have received particular attention in recent years as an important drug discovery modality. Therefore, accurate prediction of the modes of peptide binding to their target proteins will lead to accelerated peptide drug discovery. Although some difficulties remain that need to be overcome owing to the low number of peptide complexes in the PDB at present, development of a dataset for protein–peptide docking is also currently underway [30]. Although archive sets of docking solutions (Dockground [31] and ZLAB decoy set [32]) are available in the field of protein-protein docking, no such equivalent resource is available in the field of protein– peptide docking. The construction of a large-scale archive set of protein–peptide docking solutions and the development and exhaustive evaluation of methods based on these solutions are needed to accelerate peptide drug discovery as an important challenge to undertake.

## Acknowledgements

This work was supported in part by KAKENHI (grant nos. 15K16081, 18K18149, 20H04280) from the Japan Society for the Promotion of Science (JSPS) and the Mizuho Foundation for the Promotion of Sciences.

